# Structural basis of virion assembly in the *Mycobacterium tuberculosis*-infecting mycobacteriophage SWU1

**DOI:** 10.64898/2026.11.08.765204

**Authors:** Qucheng Bian, Yu Huang, Tingting Feng, Chongzheng Fan, Yuming Wei, Hongxiang Xu, Sirui Liu, Qianqian Sun, Xiaojie Zhang, Junhao Zhu, Xiuna Yang, Wei Zhang, Xinwen Chen, Zihe Rao, Siguo Liu, Jianping Xie, Haitao Yang, Yan Gao

**Author notes:** These authors contributed equally (Qucheng Bian, Yu Huang, Tingting Feng). Corresponding author. Correspondence to Siguo Liu.; Jianping Xie.; Haitao Yang.; Yan Gao.

## Abstract

The rising threat of drug-resistant tuberculosis has renewed interest in bacteriophage therapy as an urgently needed alternative. Rational engineering of these viruses, however, requires an atomic-level understanding of virion architecture. Mycobacteriophage SWU1 belongs to subcluster A2, one of the few mycobacteriophage lineages known to infect *Mycobacterium tuberculosis* (*M. tuberculosis*). Using single-particle cryogenic electron microscopy (cryo-EM), we resolved essentially the entire SWU1 virion architecture by localized reconstruction of four modules, the capsid, portal vertex, tail and baseplate, at resolutions of 3.62 Å, 3.51 Å, 3.70 Å and 3.22 Å, respectively. The T=7 capsid comprises 415 copies of an HK97-fold major capsid protein and lacks any decoration protein in density. The 12-fold portal reconciles its C5–C12 mismatch with the capsid vertex through defined, non-equivalent subunit contacts. The baseplate carries a 13-fold tail-fiber ring on a 6-fold hub, a geometry without precedent among characterized phages, established here by symmetry-free reconstruction. These findings provide the critical insights into the assembly and infection of SWU1, laying a structural foundation for the future optimization of phage-based anti-TB therapeutics.

## Introduction

Tuberculosis (TB) remains the leading cause of death from a single infectious agent^1^. The drug-resistant tuberculosis (DR-TB), particularly multidrug-resistant and rifampicin-resistant (MDR/RR-TB) strains, has complicated therapeutic interventions profoundly^2^. Although new anti-TB drugs such as bedaquiline, delamanid and pretomanid have improved treatment options, resistance against these agents is already emerging at an alarming pace^3,4^. Therefore, there is an urgent need for alternative anti-TB therapeutics.

To face the problem of antimicrobial resistance, bacteriophage (phage) therapy has gained traction as a highly promising strategy^5^. Phages are bacteria-specific viruses, and those capable of productive lytic growth possess the inherent ability to rapidly lyse and destroy target pathogens^6^. Most phages pose no threat to humans, yet billions of virions naturally traverse the epithelial cell layers and disseminate throughout the body via transcytosis each day^7^. Recently, phage therapy has demonstrated encouraging clinical outcomes across a range of bacterial infections. Successful cases have been reported for pathogens including *Pseudomonas aeruginosa* and *Staphylococcus aureus*, with some patients achieving infection clearance after conventional antibiotic regimens had failed^8–10^. The combination of bacteriophages with antibiotics has also shown positive effects in various clinical practices^11,12^. With respect to mycobacterial infections, a patient with disseminated *Mycobacterium abscessus* infection was successfully cured with a tailored phage cocktail^13^. Furthermore, recent studies on humanized mouse models have demonstrated that bacteriophage administration DS6A and D29 could enhance the clearance of *M. tuberculosis*, suggesting that phage-based approaches may hold potential for tuberculosis treatments^14^.

The mycobacterial cell wall is architecturally distinct from that of other bacteria, which consists of a thick layer formed by mycolyl-arabinogalactan-peptidoglycan (mAGP) complex^15,16^. Driven by the unique structural complexity of this cell wall, the majority of mycobacteriophages have evolved into the Siphoviridae family, featuring characteristically long, flexible, non-contractile tails and encoding large tape measure proteins (TMP)^17^. Some of the TMPs harbor muralytic domains, enabling them to function not merely as DNA transit channels but also as enzymatic effectors, that degrade the thick cell wall, thereby facilitating phage DNA delivery to the plasma membrane^18,19^. Several mycobacteriophages from cluster K and subclusters A2-A3 of cluster A are able to infect and lyse *M. tuberculosis* (Extended Data Fig. 1a)^20^. Moreover, a subset of these phages exhibits a broad host range, capable of infecting multiple Mycobacterium species^21^. Although high-resolution structures of mycobacteriophages such as Bxb1^22^ and Douge^23^ have recently been reported, their limited infectivity toward *M. tuberculosis* (Bxb1 forms plaques on *M. tuberculosis* at an efficiency of plating of only ∼10^-2^ relative to *Mycobacterium smegmatis* (*M. smegmatis*), whereas subcluster A2 phages such as D29 and L5 plate at 0.59 and 7.5, respectively^24,25^) means neither represent a phage that efficiently targets this pathogen. Thus, the architecture of a mycobacteriophage that efficiently infects *M. tuberculosis* remained unknown.

Mycobacteriophage SWU1, firstly isolated from a soil sample in Sichuan Province, China^26^, belongs to subcluster A2 and exhibits potent lytic activity against both *M. smegmatis* and *M. tuberculosis*^27,28^. The SWU1 genome is 52,474 bp in length and its gene content is highly similar to those of the well-characterized mycobacteriophage L5^28,29^. Notably, SWU1 encodes a unique protein gp39^30^, and expression of gp39 in mycobacterial cells increases cell wall permeability, thereby sensitizing the bacteria to a broad spectrum of antibiotics^30^. Despite its significant therapeutic potential, high-resolution structural information for SWU1 remains absent, hindering a mechanistic understanding of its particle assembly and host recognition. Here we report the near-complete structure of the SWU1 virion at near-atomic resolution, highlighting the distinct structure features not previously observed in characterized mycobacteriophages. This insight not only delineates how the SWU1 virion is assembled but also identifies candidate determinants of host recognition, providing a structural framework for the rational design of phage-based therapeutics against *M. tuberculosis*.

## Results

### Overall architecture of mycobacteriophage SWU1

Mycobacteriophage SWU1 was propagated in the permissive host *M. smegmatis*^26^. We verified that SWU1 could productively infect *M. tuberculosis* H37Ra. In spot-based plaque assays, SWU1 formed discrete plaques on *M. tuberculosis* H37Ra lawns across the dilution series, with plaque numbers comparable to those of mycobacteriophage D29, a well-characterized *M. tuberculosis* phage (Extended Data Fig. 1b)^14^. In growth-based lysis assays, SWU1 reduced the OD_600_ of infected H37Ra cultures to a level comparable to that of D29 (Extended Data Fig. 1c). Then the SWU1 virions were purified by cesium chloride density-gradient centrifugation for structural determination^31,32^. Cryo-EM micrographs of the purified sample revealed structurally homogeneous virions with a canonical siphovirus morphology (Fig. 1a): an icosahedral capsid (gp81) of ∼65 nm in diameter, a flexible tail of ∼120 nm (including part of the portal vertex), and a distal baseplate of ∼38 nm (Fig. 1b). The complete genome of mycobacteriophage SWU1 has been previously sequenced and well characterized (Fig. 1c)^28,29^. In accordance with the canonical genomic organization of cluster A, the structural genes are all located on the left arm of the genome^33,34^. Given the large size of the intact virion and the conformational heterogeneity introduced by its flexible tail, conventional whole-particle refinement did not yield a satisfactory reconstruction. We therefore adopted a localized reconstruction strategy, partitioning the virion into four modules that were refined independently: the head (capsid), the neck (portal and connector complex), the tail, and the baseplate (Extended Data Figs 2 and 3).

**Fig. 1:**
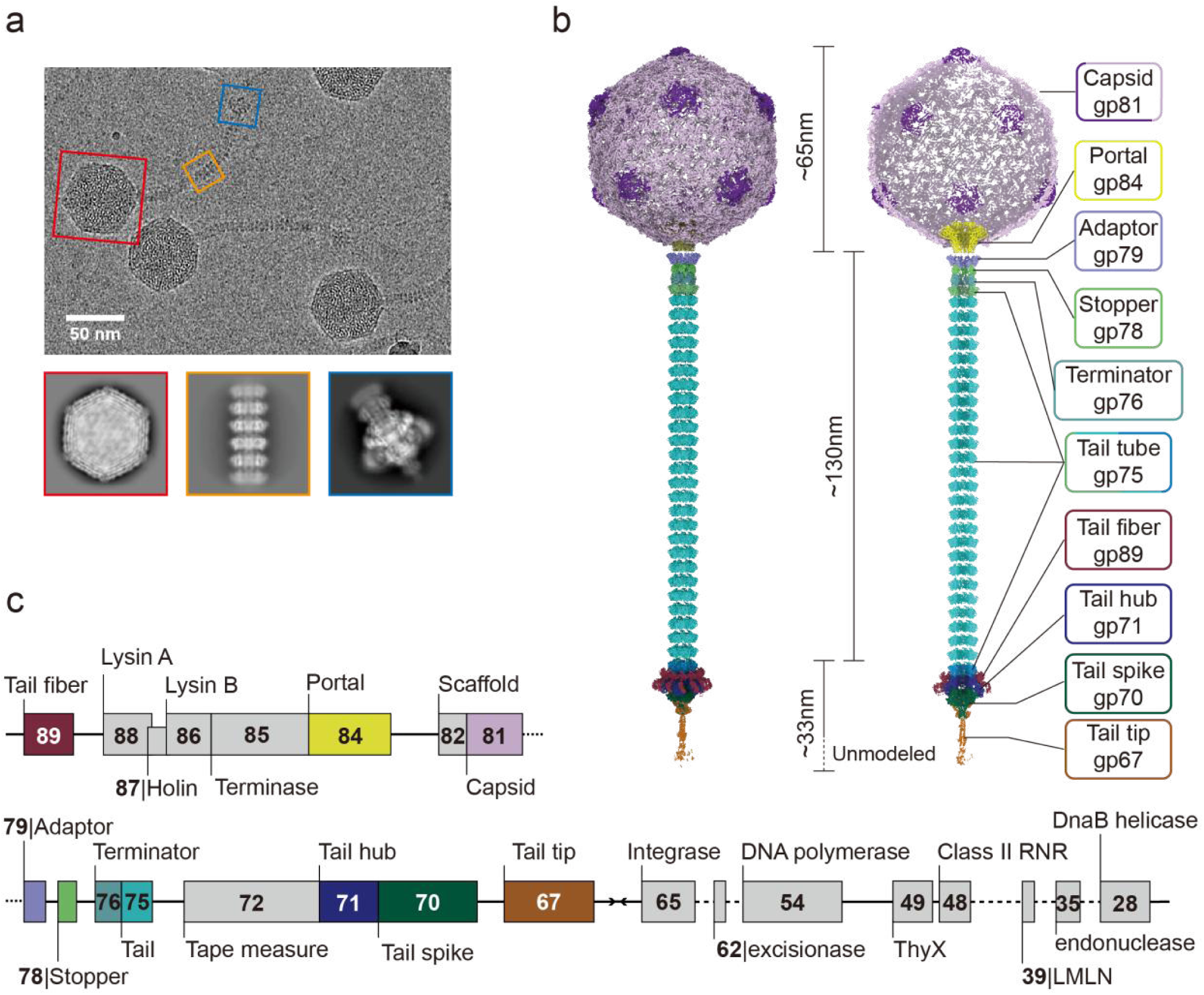
Cryo-EM structure of Mycobacteriophage SWU1. **a**, A presentative micrograph showing the overall morphology of SWU1 (scale bar: 100 nm). The capsid, tail and baseplate are highlighted in red, orange and blue boxes respectively. The corresponding 2D averages are shown below. **b**, Composite density map of SWU1 virion. The image left shows the intact volume and the right is a z-clipped map. The region indicated by the dashed line remains unmodeled due to poor map density. The structural components are colored based on the corresponding gene map in **c**. **c**, Schematic map of the viral genome. The central converging arrows indicate a switch in transcription direction. This annotation provides the gene numbering used throughout the present study. ThyX: Flavin-dependent thymidylate synthase. LMLN: Leishmanolysin-like peptidase.

Icosahedral refinement of the capsid yielded a 3.89 Å map, and local refinement of the capsid protomer improved the resolution to 3.62 Å. Particle subtraction and 3D classification isolated the portal vertex, giving a 3.51 Å C1 map of the capsid–portal complex. The adaptor, stopper and terminator together with the adjoining tail ring were refined with C6 symmetry to 3.25 Å, and straight segments of the main tail tube were refined with C6 symmetry to 3.70 Å. The baseplate complex, comprising the hub, spike, fibers and the directly attached tail tube, was reconstructed under C3 symmetry combined with symmetry expansion to a final resolution of 3.22 Å, and the proximal portion of the tail tip connected to the spike reached 3.33 Å (Extended Data Figs. 2,3 and Extended Data Table 1). Using the annotated genome sequence and these maps, we built atomic models for 10 of the 11 major structural proteins, including 482 monomers (Fig. 1b, Extended Data Fig. 4a and Extended Data Table 2). These models define the subunit composition of the virion across all four modules: the capsid is built from the major capsid protein gp81; the neck, from the portal, adaptor, stopper and terminator (gp84, gp79, gp78 and gp76), which together bridge the capsid and tail into a continuous conduit for DNA transit; the tail, from the tail tube protein gp75; and the baseplate comprises the tail hub, tail fibers, tail spike and tail tip (gp71, gp89, gp70 and gp67), the canonical adsorption apparatus of tailed phages (Fig. 1b). The tape measure protein (TMP) could not be modeled because its density within the tail tube is disrupted by conformational flexibility, and the distal tail tip and peripheral tail fibers likewise remain unmodeled owing to their intrinsic flexibility. The architecture of each module is described in the following sections.

### Assembly and intrinsic flexibility of the SWU1 capsid

The SWU1 capsid is assembled with T=7 icosahedral symmetry and contains 415 copies of the major capsid protein (MCP) arranged into 60 hexons and 11 pentons (Fig. 2a)^35^,with one icosahedral vertex replaced by the portal complex. Each MCP adopts the canonical HK97 fold^36,37^, comprising an N-terminal arm (N-arm), a periphery domain (P-domain), an extended loop (E-loop), an axial domain (A-domain) and a glycine-rich loop (G-loop) (Fig. 2b). The A-domain sits at the core of both hexons and pentons (Fig. 2c and Extended Data Fig. 5a), while the E-loop and P-domain reach out to engage neighboring MCPs (Fig. 2d). The P-domain carries a P-loop that, as in mycobacteriophage Bxb1^22^, forms trimeric turret-like protrusions at the 3-fold axis and contacts the E-loops of adjacent subunits (Fig. 2e)^38^. The result is an interlocking network that holds neighboring subunits firmly in place. The SWU1 MCP, however, lacks the C-terminal extension present in Bxb1 (Extended Data Fig. 6a).

**Fig. 2:**
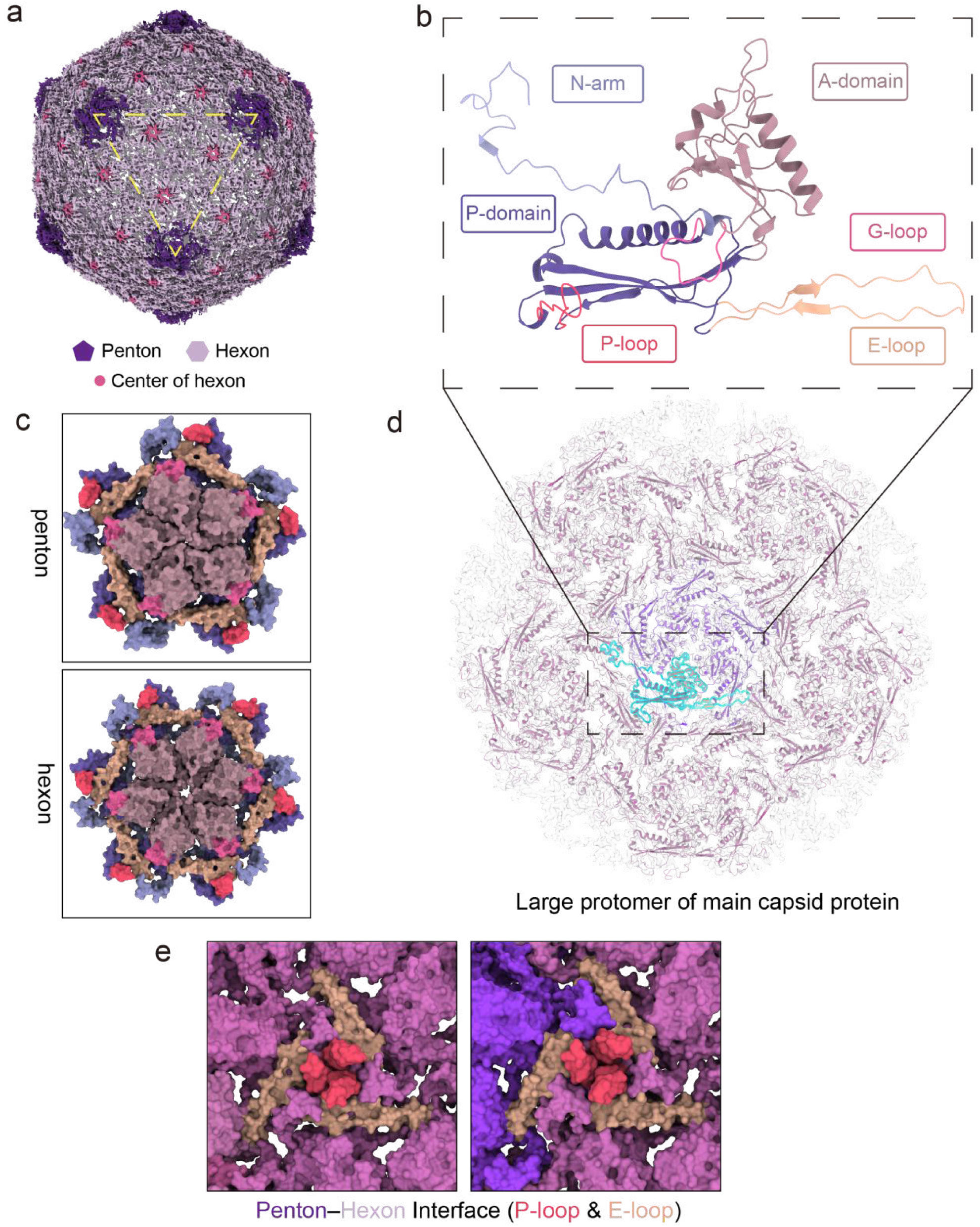
Cryo-EM structure of SWU1 capsid. **a**, Cryo-EM map of the bacteriophage capsid viewed along the icosahedral 3-fold axis of symmetry. The pentons at the vertices are colored in purple, and the hexons forming the facets are shown in light purple. The center of the hexons are highlighted in pink for visual clarity. **b**, The protein fold of a capsid subunit, colored by domains and key loops. N-arm, light purple; A-domain, mauve; P-domain, dark blue; P-loop, red; E-loop orange. **c**, Pentons and hexons showing in surface. **d**, The protein fold of the large protomer assembly. A single subunit is highlighted in cyan to illustrate its location and structural context within the complex. The dashed box corresponds to the isolated monomer detailed in **b**. **e**, Interaction interface between adjacent pentons and hexons.

Unlike the capsids of previously characterized mycobacteriophages^22,23^, the SWU1 capsid is markedly flexible, which is why the global map reaches only 3.89 Å while the protomer refines locally to 3.62 Å (Extended Data Fig. 2). We restricted all capsid reconstructions to mature, DNA-packaged particles, so the flexibility we see is not a by-product of genome release. Many HK97-fold capsids carry extra domains or decoration proteins that plug into MCP interfaces and stiffen the shell^38,39^, SWU1 has neither: no such density appears in our capsid map (Extended Data Fig. 5b), and no candidate decoration gene exists in its genome^28^. Their absence may explain why the SWU1 capsid is so flexible (Extended Data Fig. 5b). All-atom molecular dynamics simulations of the HK97 virion have likewise indicated that packaged DNA deforms the capsid shell^40^, hinting that some degree of elasticity may be built into every assembled HK97-fold capsid.

### Symmetry mismatch at the SWU1 portal vertex

The portal complex of SWU1 comprises four components—the portal, adaptor, stopper and terminator—that together bridge the capsid and the tail (Fig. 3a, b and Extended Data Fig. 5c). In tailed bacteriophages this connector is the structural hub through which the genome is packaged, retained and ejected^41–43^. In SWU1 the portal and adaptor each assemble with 12-fold symmetry while the stopper and terminator are 6-fold, the canonical 12-12-6-6 transition seen in other siphophages (Fig. 3a and Extended Data Fig. 6b)^22,23,44^. The complex replaces one capsid penton at an icosahedral vertex, so the 12-fold portal meets the 5-fold geometry of the surrounding capsid: a local C5–C12 mismatch at the capsid–portal interface (Fig. 3c). In tailed phages and herpesviruses such mismatches are generally absorbed by structural plasticity of the portal ring^45–47^; in SWU1, as described below, that plasticity takes the form of a defined set of non-equivalent subunit contacts.

**Fig. 3:**
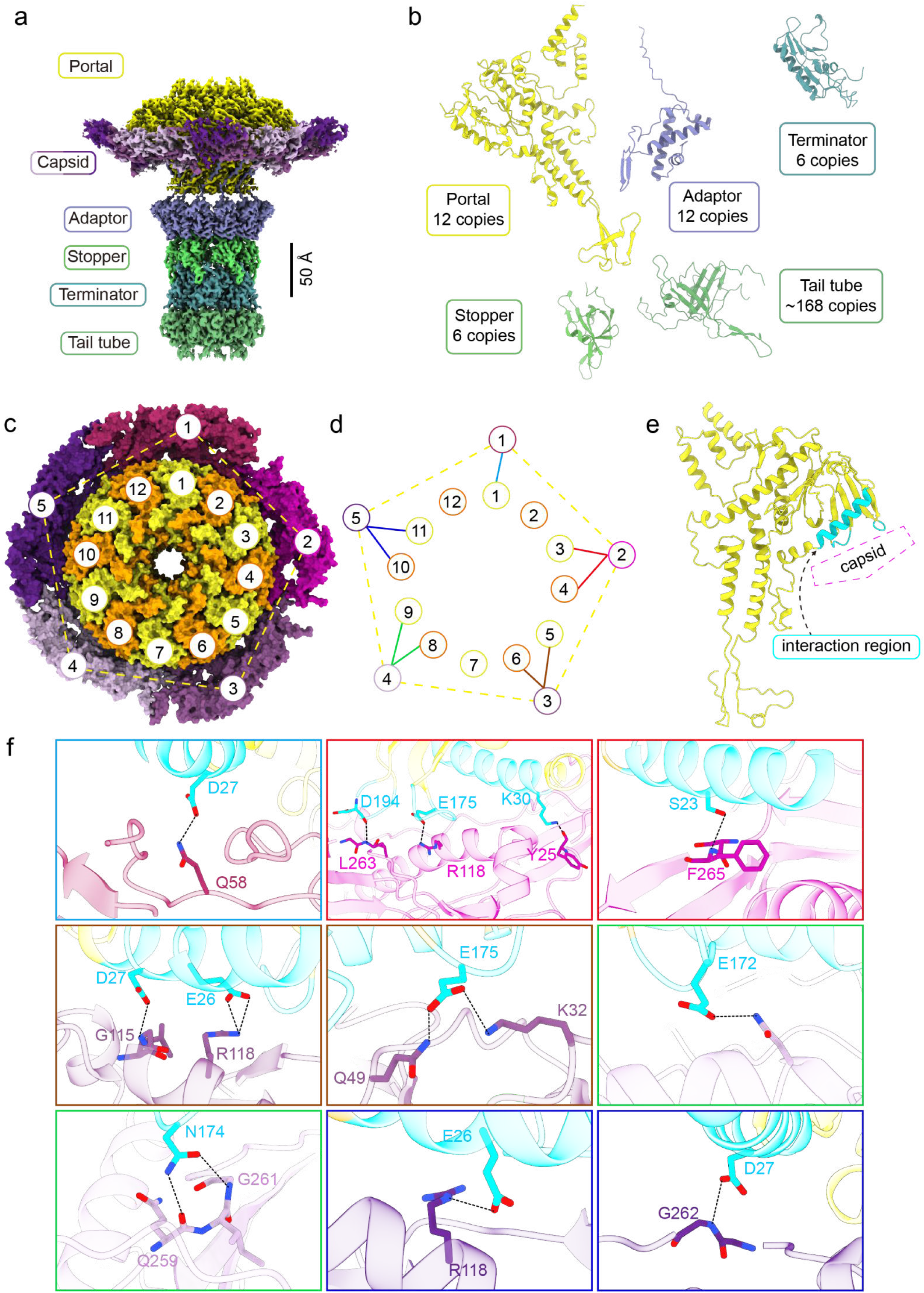
Cryo-EM structure of SWU1 portal complex. **a**, Composite density map of SWU1 portal complex. The map is colored by the indicated structural components. Components are colored individually: portal (yellow), adaptor (purple blue), stopper (green), terminator (teal blue), tail tube (light green). Scale bar, 50 Å. **b**, Cartoon diagrams of the atomic models for the subunits of the portal, adaptor, terminator, stopper, and tail tube proteins. The colors correspond to the assigned densities in **a**. **c**, Surface representations of the portal-capsid interface viewed along the central axis. Individual subunits of the 12-fold portal (yellow/orange) and the 5-fold capsid vertex (shades of purple) are numbered to show the symmetry mismatch. **d**, Schematic diagram illustrating the interaction network between the capsid and the portal. Colored lines indicate hydrogen-bonding interfaces between specific subunits. Dashed lines represent the subunit inner interactions. **e**, Cartoon diagrams showing the interaction region (highlighted in cyan) of the portal subunit with the capsid. **f**, Close-up views of the specific hydrogen-bonding interactions corresponding to the colored lines in **d**. Hydrogen bonds are denoted by red dashed lines.

To resolve the capsid–portal interface, we extracted portal-containing sub-particles from the icosahedral capsid and retained those with detectable portal signal. Each sub-particle was then symmetry-expanded five-fold about the vertex axis (C5), generating five copies that differ only in the rotational register of the 12-fold portal relative to the capsid vertex. 3D classification separated these copies, and duplicates that did not match the native register were discarded; C1 refinement of the resulting stable class gave a 3.51 Å map of the capsid–portal assembly (Extended Data Fig. 2). For the distal connector—the adaptor, stopper, terminator and the adjoining tail ring—subtracted particles were re-centered on the sub-complex and refined with C6 symmetry to 3.25 Å (Extended Data Fig. 2). Together these maps describe the portal complex in full, from the capsid shell to the proximal tail.

The resulting models reveal how the 12-fold portal reconciles itself with the 5-fold vertex. Rather than contacting the capsid equivalently, the portal subunits form a defined set of non-equivalent interfaces with the five hexons that encircle the vertex (Fig. 3c, d): hexon 1 is engaged by a single portal subunit (subunit 1), whereas each of hexons 2–5 is engaged by a pair of adjacent portal subunits (3 and 4, 5 and 6, 8 and 9, and 10 and 11). Nine of the twelve portal subunits thus contribute to the interface; the remaining three (subunits 2, 7 and 12) sit at hexon–hexon junctions without a detectable hydrogen-bonded contact. On the portal side of the capsid-portal interface, contacts are contributed by a composite surface formed by one α-helix (portal residues 14–31) and two loops (portal residues 171–175 and 193–197) (Fig. 3e and Extended Data Fig. 5d). The contact pattern is therefore discrete and geometry-adapted rather than averaged: the portal is anchored at the vertex by subunit-specific interactions, and this is how SWU1 accommodates the C5–C12 mismatch (Fig. 3f and Extended Data Fig. 4b). For instance, at the hexon 1–portal subunit 1 interface, the side-chain amide nitrogen of MCP Gln58 forms a hydrogen bond with the side-chain carboxyl oxygen of portal Asp27. Yet the hexon 2 interface with portal subunits 3 and 4 is more elaborate: the backbone amide nitrogen of MCP subunit Leu263, the guanidinium nitrogen of MCP Arg118 and the backbone carbonyl oxygen of MCP Tyr25 in hexon 2 pair, respectively, with the backbone carbonyl oxygen of portal Asp194, the side-chain carboxyl oxygen of portal Glu175 and the side-chain amino nitrogen of portal Lys30 in subunit 3—all drawn from the α-helix and loops defined above; while the backbone carbonyl oxygen of MCP Phe265 in hexon 2 donates to the side-chain hydroxyl oxygen of portal Ser23 in subunit 4 (Fig. 3f). Each hexon–portal interface presents its own hydrogen-bond set (Fig. 3d, f). Such an interface is necessarily mechanically anisotropic. In this geometry, the paired-contact subunits may carry most of the load imposed by genome packaging ^45,48^. Downstream, the adaptor maintains continuity: its C-terminal segment wraps around the portal ring and couples the 12-fold portal to the 6-fold stopper and terminator below (Extended Data Fig. 5e).

### Dynamic conformations of the SWU1 tail tube

The SWU1 tail tube is intrinsically flexible, a property documented across siphovirus tail tubes and a major obstacle to high-resolution reconstruction^49–51^. We therefore performed helical refinement on straight tube segments only, which yielded a 3.70 Å map; by construction, this map describes the straight conformation of the tube (Extended Data Fig. 3 and 4a). The tube is a helical stack of C6 rings with a rise of 45.41 Å and an inter-ring twist of 16.17° (Fig. 4a), and its lumen has an inner diameter of ∼39 Å (Fig. 4b). Running along the lumen axis is a continuous rod of density ∼19 Å in diameter that we attribute to the tape measure protein (TMP); this density is continuous but too weak to be interpreted as an atomic model, which is why the TMP is absent from the model above (Fig. 4b)^52,53^. Tail rings abutting the connector and the baseplate show higher local resolution than the central tube, consistent with the restricted motion imposed on them by the flanking assemblies (Fig. 4c, Extended Data Figs. 2, 3 and 7a, b)^54,55^.

**Fig. 4:**
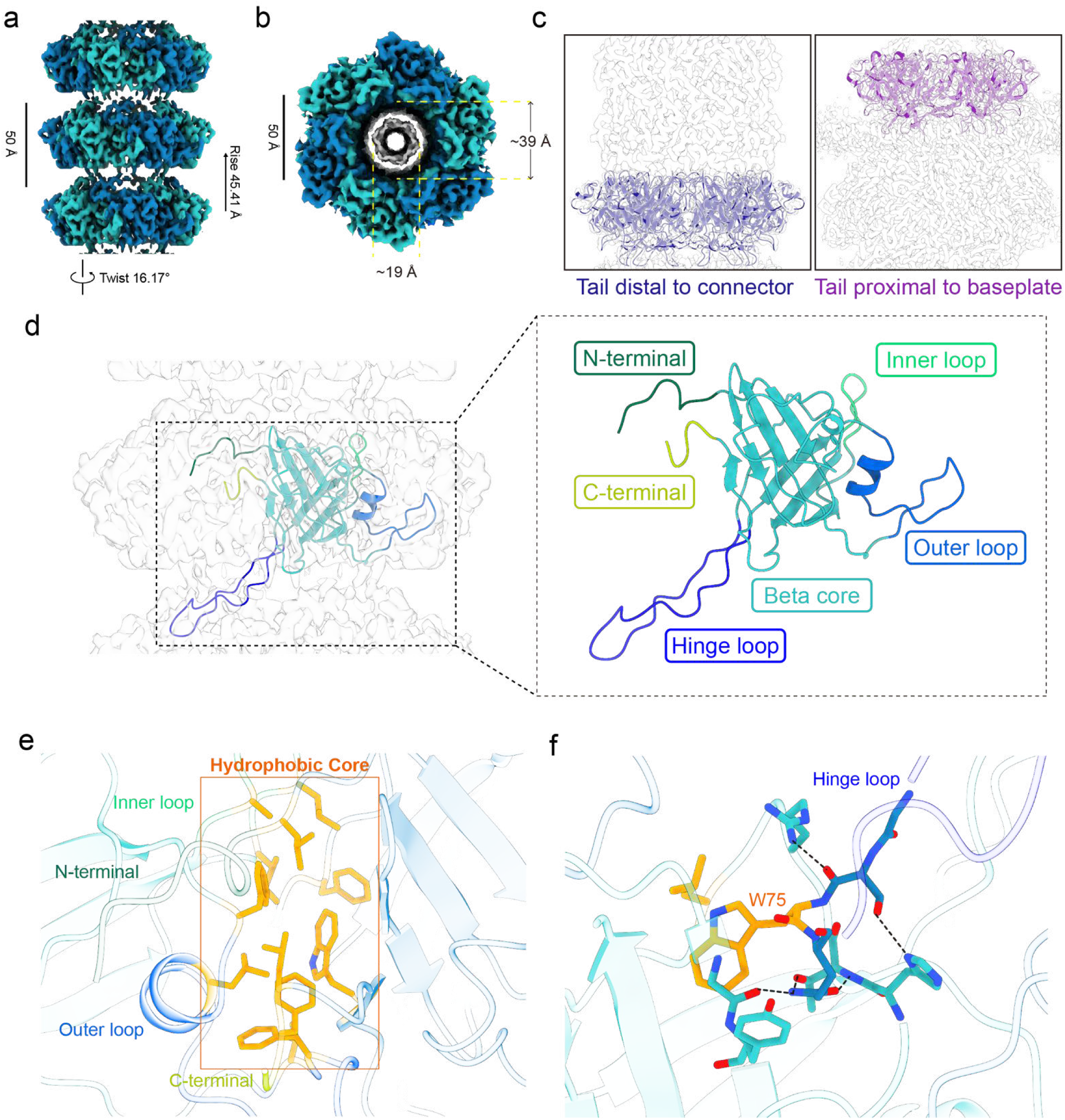
Cryo-EM structure of the SWU1 helical tail. **a**, Side view of a short segment of the density map of the helical tail. The calculated helical parameters, rise (45.41 Å) and twist (16.17°), are shown. Scale bar, 50 Å. **b**, Top view of the tail density map viewed along the helical axis. The TMP density is colored grey. Scale bar, 50 Å. **c**, Tail rings from the two terminal assemblies, shown in side view with the tail axis horizontal. Left, the ring immediately distal to the connector, colored in purplish blue; Right, the ring immediately proximal to the baseplate hub, colored in light purple. **d**, Ribbon diagram of the major tail subunit. Key domain regions are highlighted. N-terminal, dark green; inner loop, light green; C-terminal, yellow green; outer loop, royal blue; beta core, cyan; hinge loop, blue. **e**, Close-up view of the hydrophobic core (highlighted within the orange box) that stabilizes the structural arrangement of the N-terminal, C-terminal, inner loop, and outer loop regions. **f**, Detailed view of the molecular interaction network stabilizing the hinge loop. The hydrophobic core including key residue Trp75 (orange) and its associated hydrogen bonds (dashed lines) are highlighted.

Each tail tube protein (TTP, gp75) monomer is built around the conserved β-core of the canonical tail-tube fold^49,56,57^, comprising an N-terminal domain, a C-terminal domain, an outer loop and an inner loop (Fig. 4d). The same fold recurs in all three tube environments we resolved: TTP subunits from the central tail tube and from the rings adjoining the connector and the baseplate superimpose with essentially identical domain organization, and the one region that slightly differs among them is the hinge loop (Extended Data Fig. 7a, b). Within a ring, the N and C termini insert into the outer and inner loops, respectively, of neighboring subunits, burying a hydrophobic core at the subunit–subunit interface that locks the six subunits of the ring together (Fig. 4e and Extended Data Fig. 4b). Rings are stacked through the hinge loop of each TTP, which reaches into the adjacent ring; the freedom of this loop to extend, retract and deflect laterally is what allows the tube to bend (Fig. 4d, f) ^50,51^. A conserved tryptophan in the hinge loop (Trp75) plugs into the β-core of the adjacent ring, its indole side chain seated in a hydrophobic pocket and clamped by hydrogen bonds from the surrounding residues—an arrangement that explains how the rings stay attached while the tube remains bendable (Fig. 4f and Extended Data Fig. 4b) ^51,57^. At the connector and baseplate ends the same hinge loops adopt progressively more constrained conformations; this is the structural basis both of the higher local resolution noted above and of the anchoring of the tube to the terminal assemblies^53,55^.

### Structural Complexity of the SWU1 Baseplate

The baseplate complex, located at the distal extremity of the tail, serves as the critical molecular apparatus for host recognition and attachment^58,59^. Our cryo-EM maps show a bell-shaped complex built from a central C6 tail hub (gp71), a C3 tail spike (gp70) nested inside it, and an elongated tail tip (gp67) projecting downward (Fig. 5a, b). Around the hub runs a ring of tail fibers (gp89) with 13-fold. The count comes from a symmetry-free (C1) reconstruction of the tail-baseplate complex, in which the tail fibers resolve as thirteen discrete densities at thirteen distinct angular positions (Fig. 5c and Extended Data Fig. 3); the C3 refinement after re-centering to tail tip cannot generate this arrangement clearly, so the 13-fold ring is a property of the particle rather than an artifact of imposed symmetry. Fiber rings in characterized bacteriophages are almost invariably multiples of three^22,23^, with five-fold rings reported only rarely^60^; a 13-fold ring, to our knowledge, has no precedent, and it widens the known geometric range of viral assembly (Extended Data Fig. 6c). The fibers project outward and distally as the outermost elements of the virion, the geometry with the potential to act as a receptor-binding-protein (RBP) or RBP-associated protein^58,61^.

**Fig. 5:**
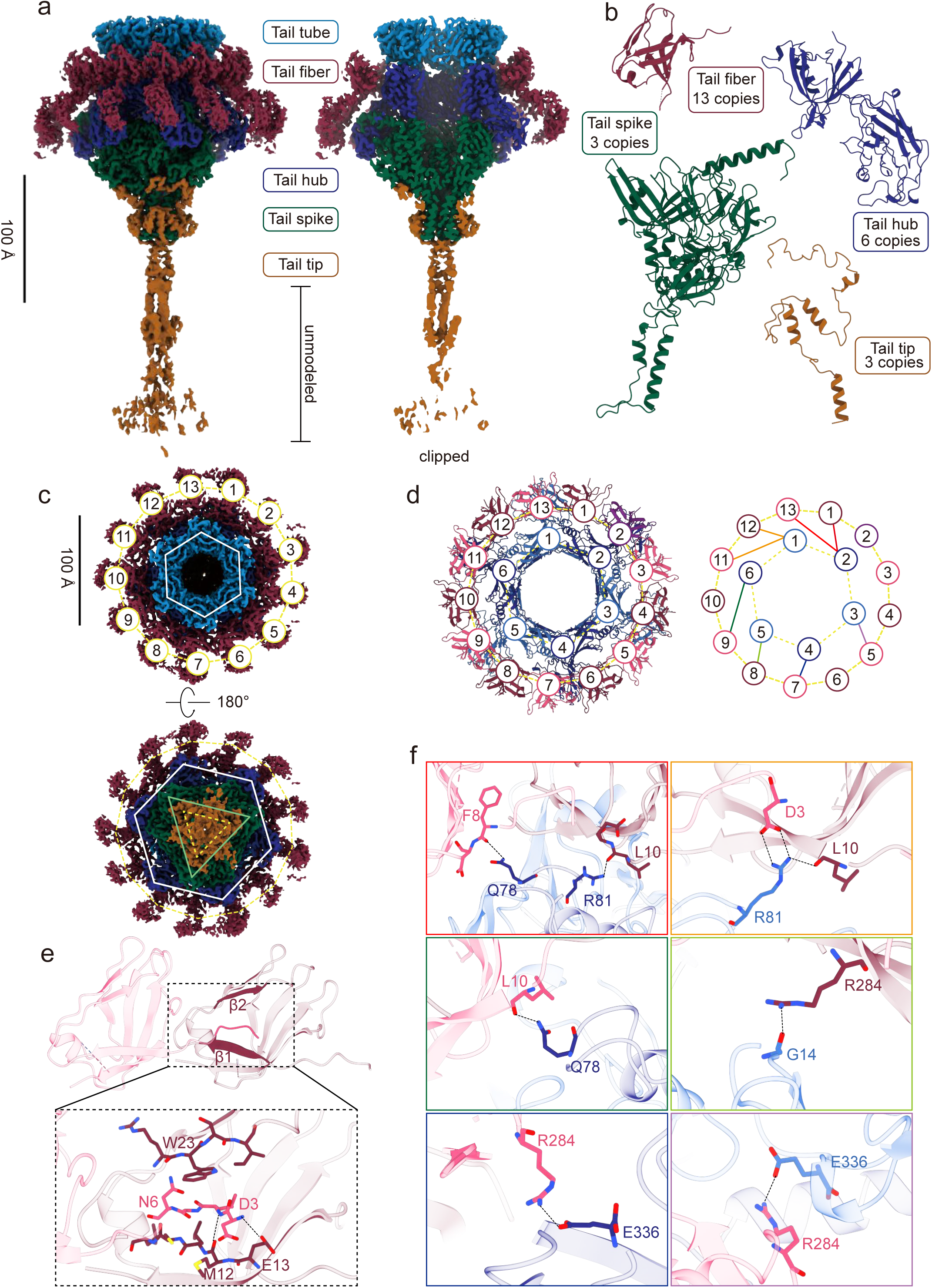
Overall architecture of the SWU1 bacteriophage baseplate complex and the molecular basis of the rare C6-C13 symmetry mismatch. **a**, Cryo-EM maps of the intact (left) and cutaway (right) baseplate complex. Components are colored individually: tail tube (light blue), tail fiber (dark red), tail hub (dark blue), tail spike (dark green), and tail tip (sienna). The highly flexible, unmodeled distal region is indicated. **b**, Atomic models and corresponding copy numbers for individual baseplate components. **c**, Top (upper) and bottom (lower) views showing the nested symmetries. The C13-symmetric tail fibers (numbered 1-13) surround the central C6-symmetric tail hub (white hexagon) and C3-symmetric tail spike (green triangle). The upper view is taken from the capsid side, looking down the tail axis toward the distal end; tail fibers are numbered 1–13 clockwise in this orientation. **d**, Molecular surface (left) and schematic topology diagram (right) of the atypical binding network between the tail hub (dark blue/steel blue) and tail fibers (dark red/purple), viewed from the capsid side. Hub monomers (1–6) and tail fibers (1–13) are numbered clockwise in this orientation. Solid lines denote defined hydrogen-bonded or salt-bridged interfaces; dashed lines denote the subunit inner interactions. **e**, Structural details of the tail fiber atomic model (pink). The inset highlights the residue interaction network within the core folded region. **f**, Close-up views of the varying interfaces between tail fibers and tail hubs. Key electrostatic interactions and dynamic salt bridges (dashed lines) are highlighted, including the specific engagement between fiber Arg284 and the extreme C-terminal Glu336 of the hub.

A 13-fold fiber ring seated on a 6-fold hub is a mismatch of a kind not previously described: 6 and 13 share no common factor, so no symmetry operation can map fiber positions onto hub positions, and no repeating hub–fiber pairing can close the ring. The hub meets it the way the portal meets the 5-fold vertex—by giving up equivalence— but the bookkeeping is different. Hub monomers 1 and 2 each grip two fibers (monomer 1: fibers 11 and 12; monomer 2: fibers 13 and 1), whereas hub monomers 3, 4, 5 and 6 each grip one (fibers 5, 7, 8 and 9, respectively; Fig. 5d). Eight of the thirteen fibers are thereby anchored to the hub through hydrogen bonds and salt bridges; the remaining five fibers 2, 3, 4, 6 and 10 form no such interactions and are held in the ring only by fiber–fiber contacts (Fig. 5d, e). Throughout the ring, adjacent fibers are linked in the same way: the N-terminal loop of one gp89 subunit (Asp3–Asn6) inserts against the β1–β2 sheet of its neighbor, forming an interface of a hydrogen-bond pair and hydrophobic side-chain stacking (Fig. 5e). The hub-fiber chemistry is no more repeating than the pairing. For example, at the hub monomer 2 interface, Arg81 of the hub donates a side-chain hydrogen bond to the backbone carbonyl of Leu10 in fiber 1, while Gln78 donates to the backbone carbonyl of Phe8 in fiber 13—two fibers held by the same monomer through two different residues. At the hub monomer 3 interface, by contrast, the contact is a salt bridge between the side-chain carboxylate of Glu336, the terminal residue of the hub (gp71), and the guanidinium group of Arg284 in fiber 5 (gp89) (Fig. 5f and Extended Data Fig. 4b). This mechanism of compensating for global symmetry breaking through local interface plasticity highlights the evolutionary adaptability of phage assembly^62,63^.

Meanwhile, the two ends of a tail fiber (gp89) behave very differently. The N-terminal region is the ordered part: it is the segment that locks each subunit into the ring (Asp3– Asn6; see above) and, in eight of the thirteen fibers, into the hub, and its density is strong enough that the AlphaFold3-predicted fold fitted it with only minor adjustments during manual rebuilding. The distal, outward-facing half is the opposite: its density is continuous but weak, the hallmark of conformational flexibility, and we left it unmodeled (Extended Data Fig. 7c). The weak density is compatible with the extended path the prediction traces, though it does not resolve it (Extended Data Fig. 7c, d). A distal arm free to swing in this way would sweep a larger area of the host surface while searching for a receptor, and may therefore raise the odds of a productive first contact^64,65^.

The tail tip adopts a C3 symmetry which N-terminal domain intricately wraps around and envelops the basal region of the tail spike, mediating a robust interconnection. Furthermore, the distally projecting needle-like terminus of the tail tip exhibits inherent structural flexibility and we also left it unmodeled (Extended Data Fig. 7e). Within the internal lumen of the baseplate, we identified an exquisite DNA control switch. The inner α-helices of the tail spike extend prominently into the central cavity, converging to form a tight physical “gate” that effectively retains the viral genomic DNA within the phage interior prior to infection (Extended Data Fig. 7f)^42,66^. This architectural design, combining a rigid anchor with a flexible probe, likely functions as the essential mechanical apparatus during the initial stages of host cell wall puncturing^67,68^.

## Discussion

In this study we present a near-complete virion architecture of mycobacteriophage SWU1: capsid, portal vertex, tail and baseplate, resolved as four modules between 3.2 and 3.7 Å. The baseplate seats a 13-fold tail-fiber ring on a 6-fold hub, a combination without precedent among characterized phages, while our portal analysis follows the framework established for Bxb1^22^ and Douge^23^, applied here to SWU1, a subcluster A2 phage that infects *M. tuberculosis*. These structures are architectural, not mechanistic. Host recognition, for example, is inferred from baseplate geometry and genomic context rather than observed. SWU1 is therefore offered as a structural platform for that next step, and as a model for the subcluster A2 phages that infect *M. tuberculosis*. SWU1 differs substantially from the previously well-characterized Bxb1; despite their common cluster A origin, comparative genomic analyses reveal profound divergence between SWU1 (subcluster A2) and Bxb1 (subcluster A1): only 38 proteins share 0.3–0.8 similarity, with most exhibiting relatively low similarity around 0.5. Their lysogeny and DNA-metabolism strategies also differ. Bxb1 integrates through a single well-characterized serine integrase–directionality factor pair (Bxb1 gp35 and gp47) acting at attP/attB^69^. SWU1 carries an equivalent cassette (integrase gp65, excisionase gp62, attP) but adds homologous-recombination candidates such as a single-strand annealing protein (gp51), endonuclease VII (SWU1 gp35) and a RecB-family exonuclease (gp23)^28^. This expanded enzymatic toolkit suggests that SWU1 possesses a superior capacity for DNA repair, replication restart, and genomic mosaicism. Cross-genus homologies point the same way: 32 SWU1 genes match *Segniliparus rugosus* (*S. rugosus*) ATCC BAA-974 and 54 find their closest homologs in *Mycobacterium rhodesiae* (*M. rhodesiae*) JS60, spanning structural, packaging and DNA-metabolism genes—a distribution interpreted as the remnants of two SWU1-like prophages in those bacterial genomes, and consistent with modular exchange and cross-genus host switching in this lineage (Extended Data Fig. 8). Tracing that history properly will take structures of more A2 phages.

SWU1 also carries accessory genes with no counterpart in the virion. The best characterized is gp39^30^. Located within an insertion fragment absent in its close relative L5, gp39 is highly conserved in only 6 subsets of phages across subclusters A2, A12 and A16, which makes it one of the most notable differences between SWU1 and its relatives. Expressed in *M. smegmatis*, gp39 compromises cell-wall integrity, raising permeability and sensitizing the bacterium to antibiotics and environmental stress^30^. Alongside this, the protein increases cellular permeability and sensitizes the bacterium to antibiotics and environmental stresses. Consequently, gp39 probably serves as a key driver for host lysis and the subsequent release and spread of progeny phages. Moving forward, structural characterization of this unique protein will be essential to unravel its precise mechanism of action within the broader context of the SWU1 infection cycle. Extending the current static structural approach to *in situ* structural analyses at the cellular level will be critical for capturing the dynamic conformational changes of gp39 throughout infection, and represents a promising direction for future investigation.

The question that which SWU1 protein actually grabs the receptor is still open for SWU1-like phages, and our structures do not close it. gp89 remains a candidate: it occupies the outermost position on the virion, its gene lies beside lysinA and lysinB in a putative infection/lysis cluster^70,71^, and it shares synteny with Bxb1 gp4^22,24^. The host range of mycobacteriophages is frequently dictated by specific surface glycolipids. For instance, phages BPs and Muddy strictly require trehalose polyphleates (TPPs) for adsorption, and TPP loss leads to resistance^72^. However, a single amino acid substitution in their tail spike can confer TPP-independent infectivity^72^. Building on this paradigm, the structural elucidation of the SWU1 adsorption interface, particularly gp89 and associated tail components, presents direct targets for structure-guided engineering. Targeted mutagenesis, domain swapping, or modular redesign of these RBPs could systematically alter receptor dependency, rationally expand the phage host range, and preemptively mitigate the risk of receptor-mutation-induced bacterial resistance.

Finally, the SWU1 virion structure, featuring its unique baseplate, establishes a molecular basis for assembly and host attachment, while highlighting the need for further investigation into its dynamic interplay with the mycobacterial cell wall. Two parts of this structure could actually be engineered. First, tail-associated receptor-binding proteins, such as gp89 mentioned above, represent direct and tractable engineering targets. Second, virion glycosylation provides another potential engineering strategy for therapeutic applications. Several mycobacteriophages carry O-linked glycans on capsid and tail tube proteins, mediated by phage-encoded glycosyltransferases^73^. These glycans can modulate antibody recognition and reduce the production of neutralizing antibodies. Thus, engineering phage glycosylation may help optimize immunogenicity, *in vivo* persistence, and repeated dosing potential in phage therapy for mycobacterial infections.

## Methods

### Phage amplification and isolation

Host *M.smegmatis* mc^2^155 is grown in 2YT liquid media to late-log phase. The cells were harvested from the culture by centrifugation at 3,500×g. The pellet was resuspended in PBS containing SWU1 (10^5^ PFU) and 2 mM CaCl_2_ and followed by incubation for 15 min. The suspension was then mixed with molten Top Agar (2YT, 100 µg/mL carbenicillin, 0.7% w/v BactoAgar, 2 mM CaCl_2_) and distributed to 2YT agar plates. After solidification, the plates were incubated at 37℃ until clustered plaques appeared. Phages were eluted by adding PBS to the plates and incubating overnight by 4℃. The eluate was clarified using a 0.22 µm filter and subsequently precipitated with 10% PEG 8000/ 1 M NaCl overnight. The pellet was collected by centrifugation at 3500×*g*, resuspended in TBS (20mM Tris-HCl, pH 8.0) and treated with DNase I at 37℃ for 30 min. Purified phages was isolated by CsCl density gradient ultracentrifugation (100,000×*g*, 4 h) using a step gradient (800, 700, 650, 600, 550, 500, 400, 300 g/L). The visible phage band was extracted and dialyzed against PBS.

### Verification of SWU1 lytic activity against *M. tuberculosis* H37Ra

The ability of SWU1 to infect and form plaques on *M. tuberculosis* H37Ra (hereafter H37Ra) was assessed by a spot-based plaque assay as previously described^74^. H37Ra was cultured in Middlebrook 7H9 broth supplemented with 10% (v/v) OADC, 0.05% (v/v) Tween 80, and 0.2% (v/v) glycerol at 37 °C with constant agitation of OD_600_ to 1. A 25 mL aliquot of the culture was harvested by centrifugation and washed twice with MP buffer (68 mM NaCl, 10 mM Tris-HCl, 10 mM MgSO_4_, 10 mM CaCl_2_). The washed cells were resuspended in MP buffer, mixed with molten top agar (Middlebrook 7H10, 0.7% w/v agar), and overlaid onto agar plates to form bacterial lawns. For the phage, a SWU1 lysate with a titer of ∼10^8^ PFU/mL was used as the starting material: 200 µL of the lysate was added to 800 µL of MP buffer, from which 10-fold serial dilutions were prepared. 3 µL of each dilution was spotted onto the dried lawns. The same dilution series and spotting procedure were applied to mycobacteriophage D29 at a comparable starting titer as a positive control. Plaques were enumerated after incubation at 37 °C for 7 days.

For the growth-based lysis assay, H37Ra was cultured in Middlebrook 7H9 broth as described above to mid-logarithmic phase. Cells were harvested by centrifugation and washed twice with MP buffer. The washed bacteria were resuspended in 100 mL of Middlebrook 7H9 broth containing 10% (v/v) OADC, 0.02% (v/v) Tween 80, and 0.2% (v/v) glycerol and adjusted to an OD_600_ of 0.3. SWU1 filtrate was added at a multiplicity of infection (MOI) of 10, followed by incubation at 37 °C with continuous shaking. Mycobacteriophage D29 was included as a positive control at the same MOI, and uninfected cultures served as the negative control. OD_600_ was measured every 24 h post-infection for 7 days. Experiments were performed in three independent biological replicates.

### Cryo-EM sample preparation and data collection

In total, 3 µL of purified phages sample (at a concentration of approximately 10^10^ PFU/mL) was applied onto a glow-discharged graphene oxide-coated, 300-mesh R1.2/1.3 Cu grid (GOQ300R1213CU50). The grid was then blotted for 3.0 s with a blot force of −1 maintained at 4℃ and 100% humidity and plunge-frozen in liquid ethane using a Vitrobot Mark IV (Thermo Fisher Scientific). Cryo-EM data collection was performed on a 300keV Titan Krios electron microscope (Thermo Fisher Scientific, USA) and with a K3 direct electron detector (Gatan, USA). Images were recorded in at 18000 × magnification using the super-resolution mode, with the final pixel size binned to 1.32 Å/pixel. A total of 12,663 movies were collected automatically using SerialEM 4.1.0beta^75^ with a total accumulated dose of 60 e^-^/Å^2^, fractionated into 40 movie frames over an exposure time of 3.4 s and with a defocus range of −1.0 to −2.0 µm.

### Cryo-EM data processing

The movie stacks were motion-corrected and dose-weighted by MotionCor2 1.6.4^76^. The contrast transfer functions and the following reconstruction and refinement were all performed in Cryosparc 4.6.1^77^.

For the capsid, several particles were manual picked for template. Then we processed template picking and 216,930 particles were picked and extracted in 640 pixels. After *ab*-initio reconstruction (I symmetry) and hetero refinement, a total of 192,913 particles were selected. Then we performed homogeneous refinement and non-uniform refinement (relaxed, maximization) and yielded a 3.89 Å map. Particle subtraction was then performed and the capsid vertex region was extracted. Local resolution of the capsid protomer was improved to 3.62 Å via 3D classification and local refinement.

To resolve the capsid-portal complex, capsid particles were subjected to symmetry expansion (I symmetry). After particle subtraction, local refinement and 3D classification, the signal corresponding to the unique portal-containing vertex was isolated. After removing duplicates, the portal region sub-particles were re-extracted in 220 pixels and yielding a dataset of 192,618 particles. Then we performed 2D classification and picked 126,666 particles. To address the C5-C12 mismatch between the portal and capsid, the particles underwent symmetry expansion (C5) and 3D classification. The duplicate particles were removed and finally a dataset of 113,142 particles was yielded. Local refinement was processed and finally 3.51 Å resolution of the capsid-portal complex was achieved (Extended Data Fig. 2).

For the connector-tail complex, the portal region sub-particle dataset was re-centered to the connector region and extracted in 220 pixels. After 2D classification, a total of 88,817 particles were selected. The particles were then re-extracted in 280 pixels and were reconstructed with C6 symmetry, reaching a resolution of 3.25 Å (Extended Data Fig. 2).

The tail tube fibers were picked using filament tracer (Diameter: 140 Å; Separation: 42 Å). The particles were extracted in 240 pixels and followed by 2D classification to select straight segments. A total of 157,317 particles were processed using helical refinement, resulting a map of 3.70 Å resolution (Extended Data Fig. 3).

The original baseplate complex 2D template was isolated from 2D classification results of tail tube fiber. After 2D classification, 218,032 particles were selected and re-extracted in 240 pixels. After *ab*-initio reconstruction and hetero refinement, a total of 184,298 particles were selected. To address the C13 tail fiber, homogeneous refinement (C3 symmetry) was processed and followed by symmetry expansion (C3) and 3D classification. After removing duplicate particles, a dataset of 183,244 particles was yielded. Followed by local refinement with C1, the peripheral 13 fiber features were resolved, and the global resolution of the baseplate reached 3.22 Å. The tail tip structure was obtained by re-centering the baseplate particles to the tip region and obtained a dataset of 175,782 particles. While the baseplate-tip interface reached 3.33 Å resolution with C3 symmetry, the distal end of the tip was resolved to only ∼6 Å due to flexibility. All reported resolutions are based on the gold-standard Fourier Shell Correlation (FSC) = 0.143 criterion ((Extended Data Fig. 3)^78,79^.

### Model building and refinement

All atomic models of SWU1 subunits were initially predicted by AlphaFold3^80^ and were rigidly docked into the cryo-EM density maps using UCSF ChimeraX 1.9^81,82^. The unfitted regions were manually built and adjusted using Coot 1.2^83^ and automated real-space refinement was performed by using Phenix 1.21.2^84^. The data validation are summarized in Extended Data Table 1. All structural figures were prepared using UCSF ChimeraX 1.9^81,82^.

---

**Extended Data Fig. 1:**
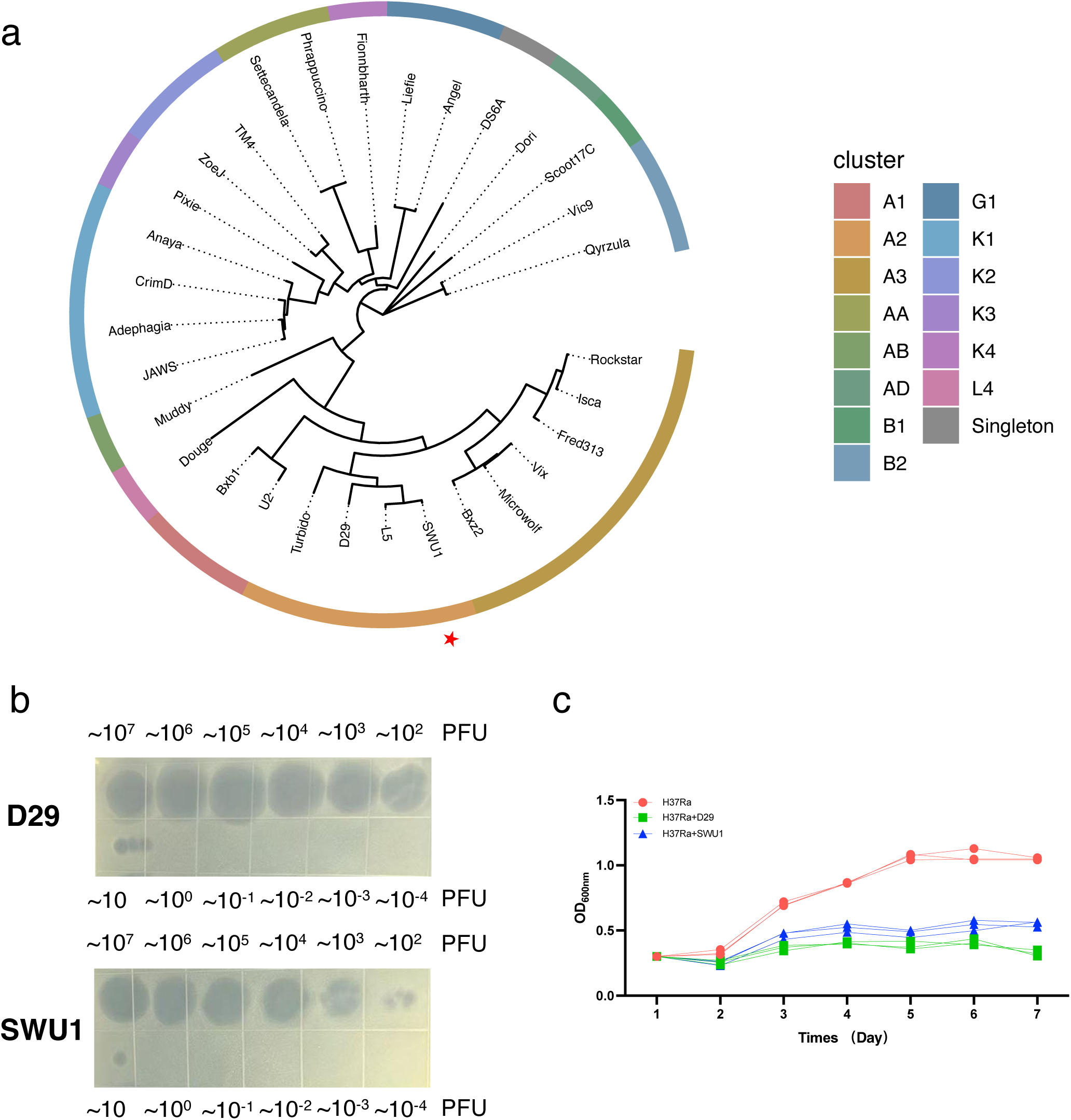
Whole-genome phylogenetic analysis and lytic activity of SWU1. **a**, Whole-genome phylogenetic analysis of SWU1 and related mycobacteriophages. Complete genome sequences of the selected phages were aligned using MAFFT, and a phylogenetic tree was subsequently constructed from the whole-genome alignment using FastTree and visualized with ggtree. Different colors in the outer ring indicate the corresponding phage clusters or subclusters, with related subclusters within the same major cluster represented by similar color schemes. The position of phage SWU1 is indicated by a red star. **b**, Spot-based plaque assay. Serial dilutions of SWU1 (starting titer ∼10^7^ PFU/mL) and of D29 were spotted onto H37Ra lawns. PFU are labelled above each row. **b**, Spot-based plaque assay of SWU1 and positive control D29. Dilution factors are labelled along each row. Both phages formed discrete plaques. **c**, Growth-based lysis assay. H37Ra cultures (OD_600_ = 0.3) were infected with SWU1 or with D29 at a multiplicity of infection (MOI) of 10, or left uninfected as a negative control, and OD_600_ was measured every 24 h for 7 days post-infection. Data are from n=3 independent biological replicates.

**Extended Data Fig. 2:**
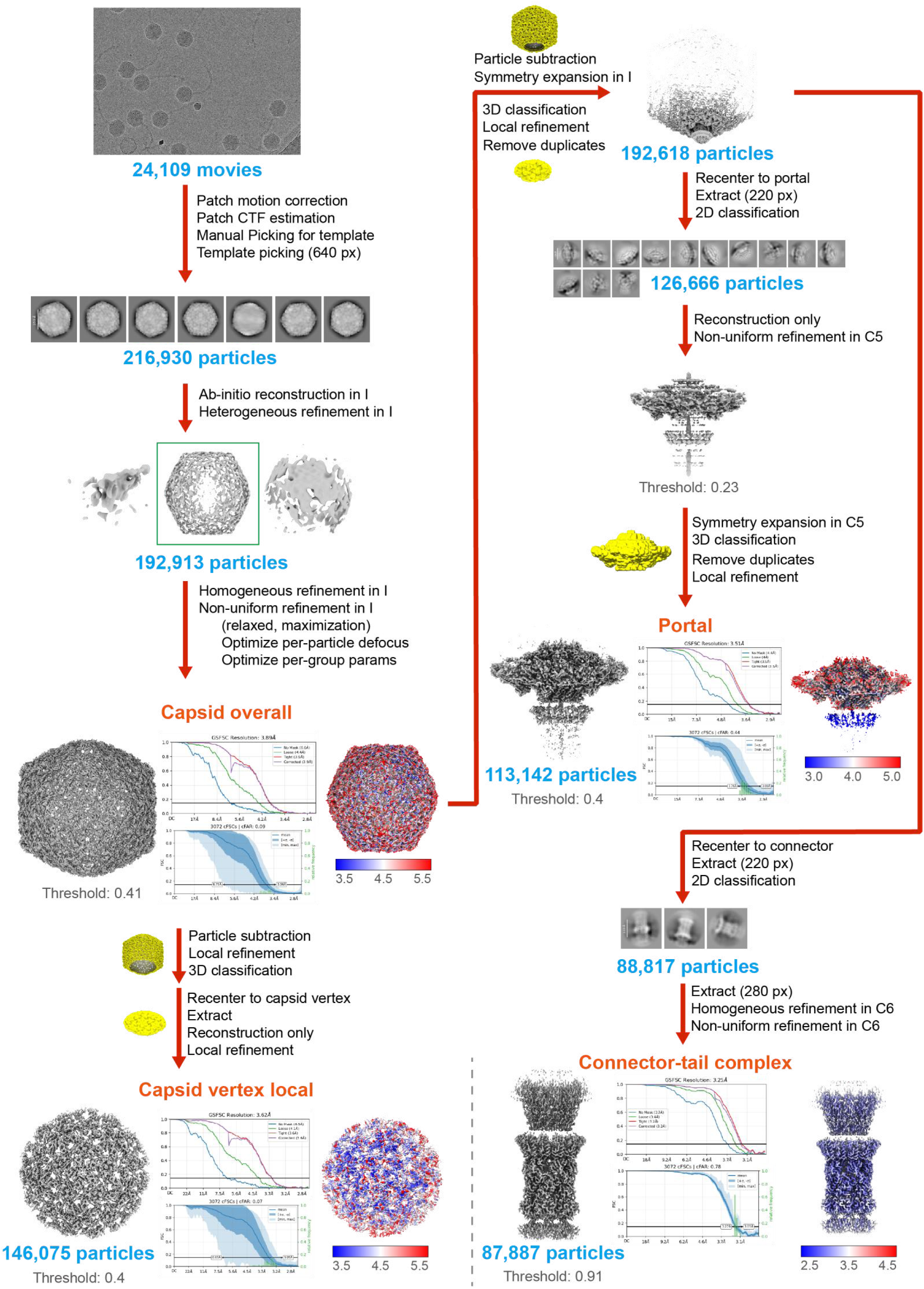
Cryo-EM data processing workflow, the GSFSC curves and the 3D FSC diagrams for SWU1 capsid overall, capsid local, portal and connector-tail complex.

**Extended Data Fig. 3:**
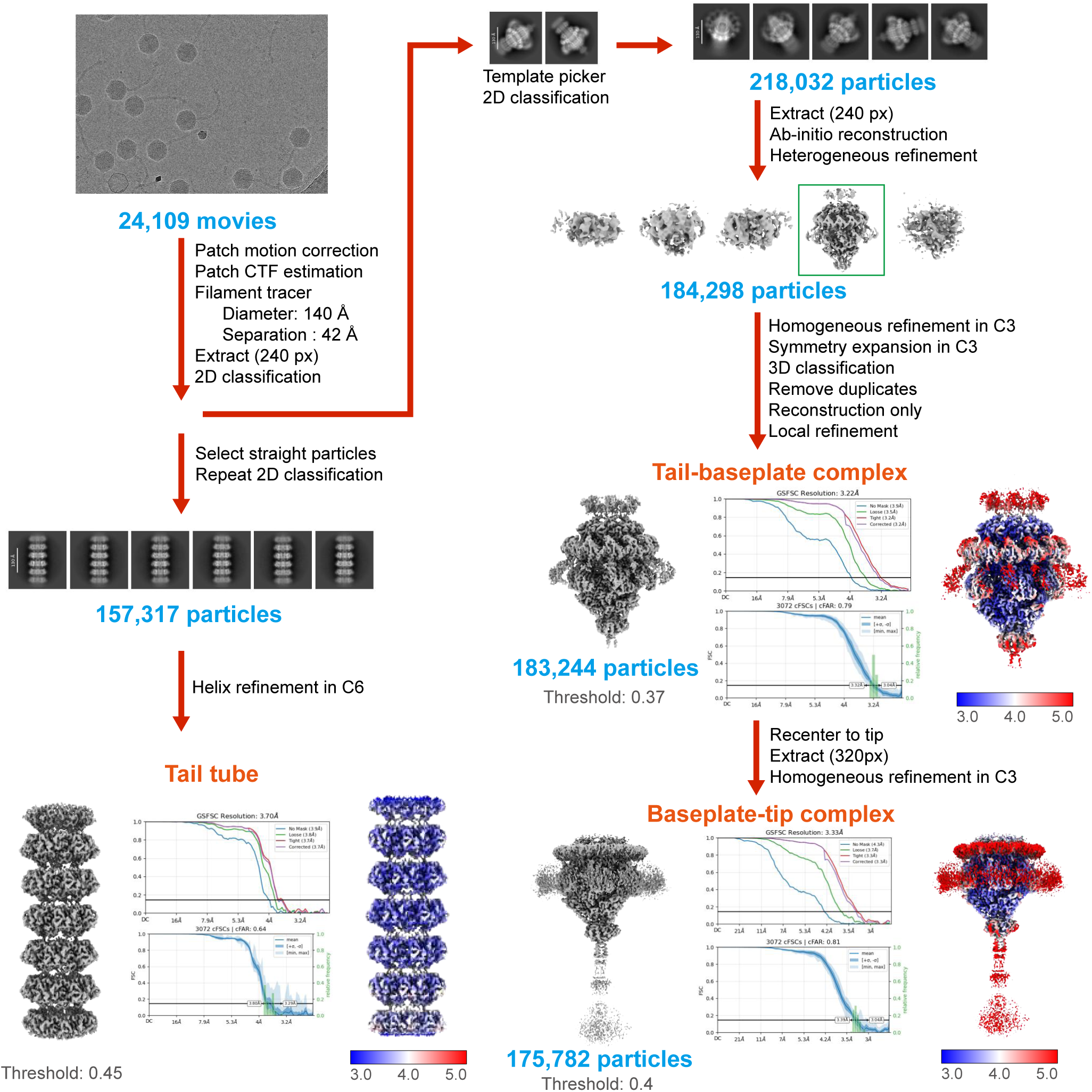
Cryo-EM data processing workflow, the GSFSC curves and the 3D FSC diagrams for SWU1 tail tube, tail-baseplate complex and baseplate-tip complex.

**Extended Data Fig. 4:**
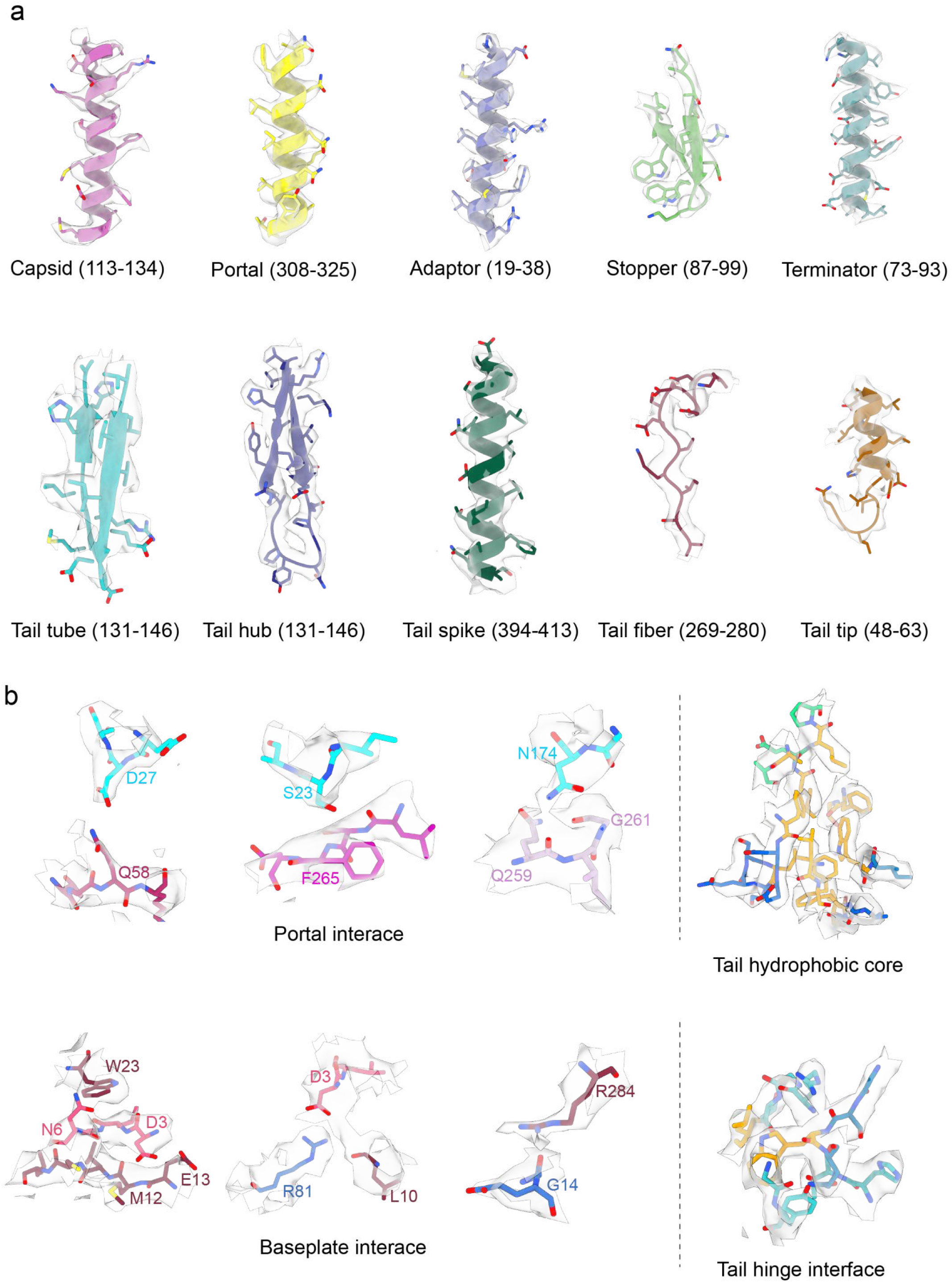
Density quality assessment of SWU1 subunits and interfaces. **a**, Representative cryo-EM densities superimposed on atomic models for polypeptides from each SWU1 subunit. **b**, Representative interaction interfaces across different structural modules, showing map densities and modeled interacting residues.

**Extended Data Fig. 5:**
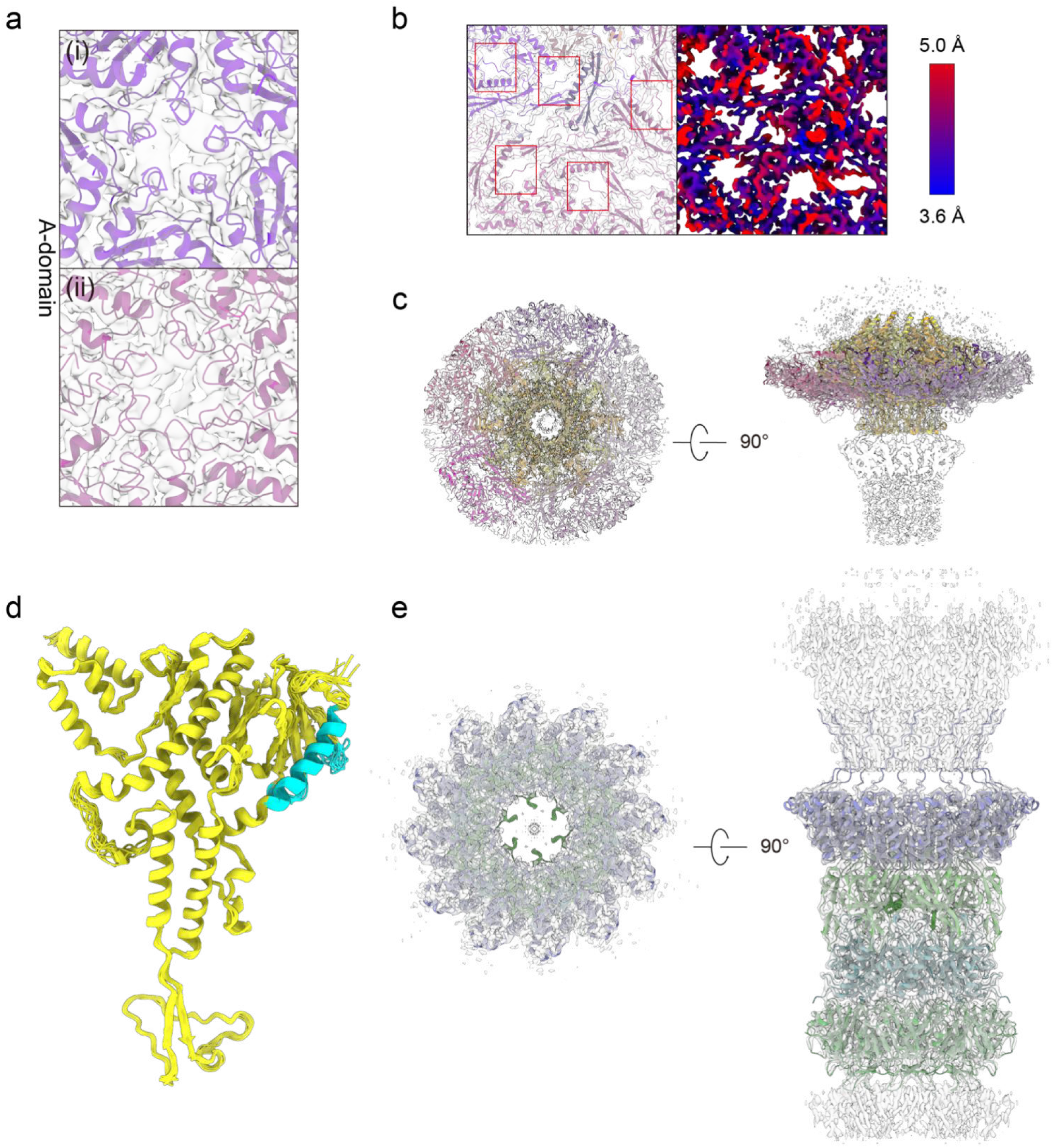
Structural details of SWU1 capsid and associated assembly complexes. **a**, Close-up view of the capsid A-domain. The pentons are colored in purple and the hexons are colored in light purple. **b**, Local resolution map of the capsid showing localized flexibility (blue, high resolution; red, low resolution). **c**, Cryo-EM map and cartoon diagram of the portal complex. **d**, Structural alignment of all portal subunits. The light blue helix marks the interface that interacts with the capsid. **e**, Cryo-EM map and cartoon diagram of the connector-tail complex.

**Extended Data Fig. 6:**
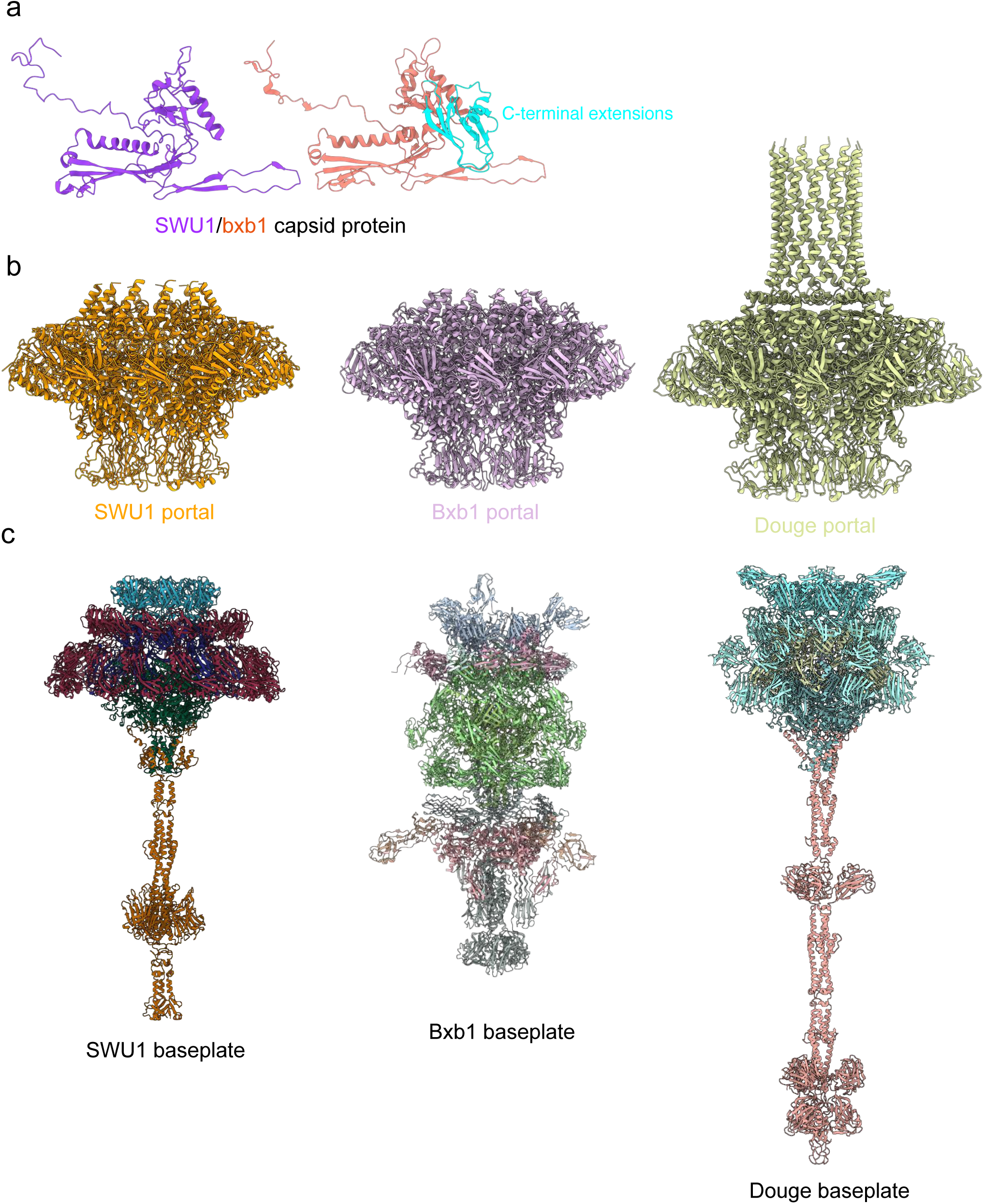
Structural comparisons of phage components among SWU1, Bxb1, and Douge. **a**, Structural comparison of the main capsid subunits between SWU1 and Bxb1 (PDB 9D9X)^22^. **b**, Comparison of the portal complexes from SWU1, Bxb1 (PDB 9D9W)^22^, and Douge (PDB 8ZDJ)^23^. The C-termini of the SWU1 and Bxb1 portals are absent, whereas the Douge portal exhibits a completely resolved C-terminus. **c**, Structural comparison of the baseplate architectures among the three phages. Bxb1 baseplate (PDB 9D93)^22^, Douge baseplate (PDB 8ZDQ)^23^.

**Extended Data Fig. 7:**
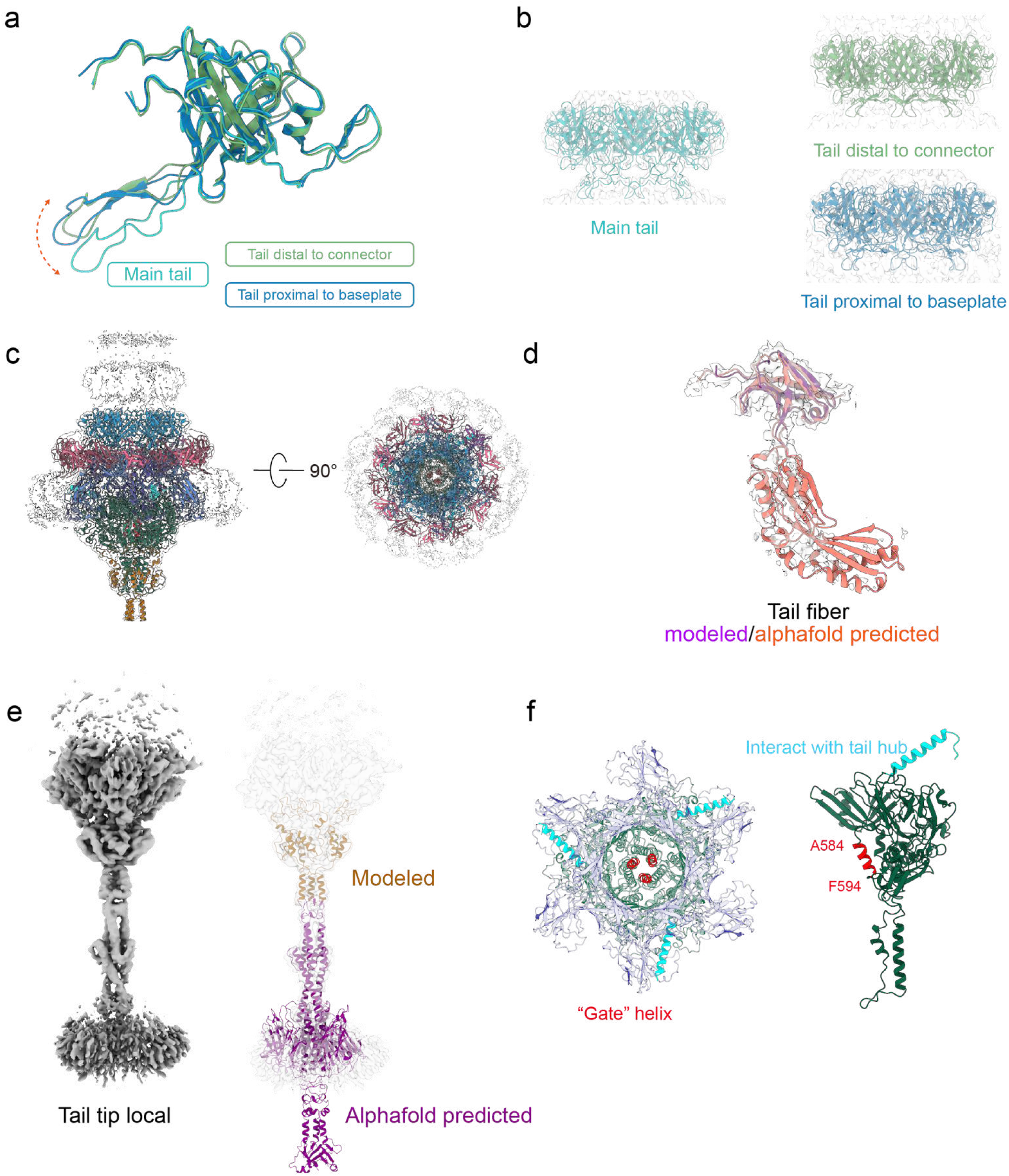
Structural insights into SWU1 tail assembly components. **a**, Superposition of major tail subunits from the main tail, tail distal to connector, and tail proximal to baseplate. **b**, Detailed views of major tail subunit densities and modeled fits for different regions shown in **a**. **c**, Structure and cryo-EM assembly map of the baseplate, with the outer region of tail fiber unmodeled. **d**, Cryo-EM density and model comparisons for the modeled and predicted tail fiber structures. **e**, Cryo-EM density and structural model comparisons for the modeled and predicted tail tip. **f**, Structural view of the phage base hub and base spike. The “gate” helices (red) (residues Ala584–Phe594) regulates the DNA ejection. The interaction region with tail hub is colored cyan.

**Extended Data Fig. 8:**
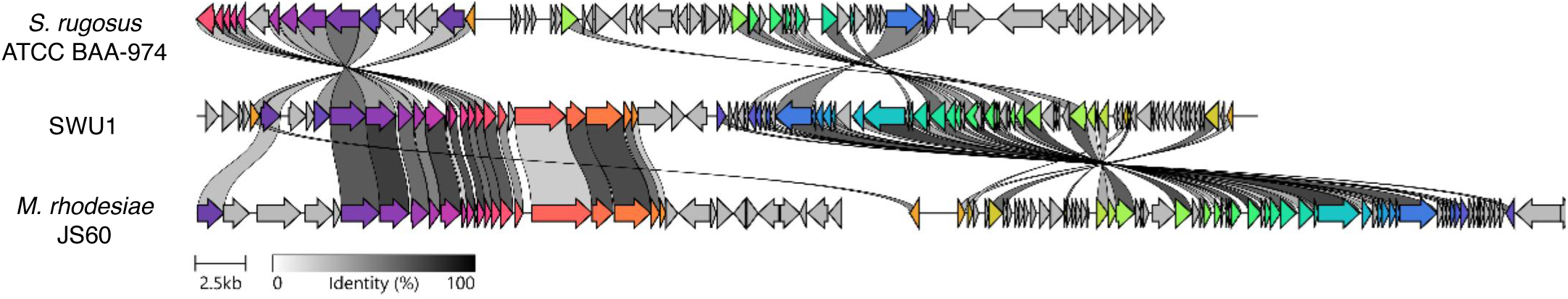
Synteny analysis of phage SWU1 and SWU1-like genomic regions in *S. rugosus* ATCC BAA-974 and *M. rhodesiae* JS60. Clinker was used to compare gene synteny among the complete SWU1 genome (NC_017973:1–52,474), two genomic scaffolds of *M. rhodesiae* JS60 (AGIQ01000018:1–31,891 and AGIQ01000007:1–33,183), and a genomic region of *S. rugosus* ATCC BAA-974 (KI391953:1,407,903–1,455,770). Arrows indicate predicted coding sequences, homologous genes are connected by shaded regions, with shading intensity indicating amino acid sequence identity. Synteny analysis revealed that close homologs of SWU1 genes in *S. rugosus* ATCC BAA-974 and *M. rhodesiae* JS60 are largely clustered in the bacterial genome and retain a certain degree of gene arrangement synteny, rather than being randomly dispersed throughout the entire genome. Therefore, these regions may represent remnants left after the integration and subsequent degeneration of SWU1-like phage genomes into the host chromosome.

**Extended Data Table 1:** Cryo-EM data collection, refinement and validation statistics.

|  | SWU1<br>capsid<br>vertex local | SWU1<br>capsid-portal<br>complex | SWU1<br>tail tube | SWU1<br>baseplate-tip<br>complex | SWU1<br>tail-baseplate<br>complex | SWU1<br>connector<br>complex | SWU1<br>capsid<br>overall | SWU1<br>tail tip<br>local |
|---|---|---|---|---|---|---|---|---|
| **PDB entry** | 43OQ | 43OP | 43OO | 43ON | 43OM | 22GZ |  |  |
| **EMDB entry** | EMD-82039 | EMD-82038 | EMD-82037 | EMD-82036 | EMD-82035 | EMD-68268 | EMD-82630 | EMD-83752 |
| **Data collection and processing** |  |  |  |  |  |  |  |  |
| Magnification | 18,000 | 18,000 | 18,000 | 18,000 | 18,000 | 18,000 | 18,000 | 18,000 |
| Voltage (keV) | 300 | 300 | 300 | 300 | 300 | 300 | 300 | 300 |
| Electron exposure (e<sup>–</sup>/Å<sup>2</sup>) | 60.00 | 60.00 | 60.00 | 60.00 | 60.00 | 60.00 | 60.00 | 60.00 |
| Defocus range (μm) | -1.2 to -2.5 | -1.2 to -2.5 | -1.2 to -2.5 | -1.2 to -2.5 | -1.2 to -2.5 | -1.2 to -2.5 | -1.2 to 2.5 | -1.2 to -2.5 |
| Pixel size (Å) | 1.32 | 1.32 | 1.32 | 1.32 | 1.32 | 1.32 | 1.32 | 1.32 |
| Symmetry imposed | C1 | C1 | C6 | C3 | C1 | C6 | I | C3 |
| Initial particle images (no.) | 216,930 | 216,930 | 4,298,352 | 1,894,029 | 1,894,029 | 2,297,342 | 216,930 | 1,894,029 |
| Final particle images (no.) | 146,075 | 113,142 | 157,317 | 175,782 | 183,244 | 87,887 | 192,913 | 175,782 |
| Map global resolution (Å) | 3.62 | 3.51 | 3.70 | 3.33 | 3.22 | 3.25 | 3.89 | 3.68 |
| Global resolution FSC threshold | 0.143 | 0.143 | 0.143 | 0.143 | 0.143 | 0.143 | 0.143 | 0.143 |
| Map sharpening *B* factor (Å<sup>2</sup>) | -89.5 | -100.7 | -177.4 | -121.8 | -100.3 | -128.2 | -79.6 | -109.5 |
| Map local resolution range (Å) | 2.9-10.0 | 3.3-10.0 | 3.0-10.0 | 2.9-10.0 | 2.9-10.0 | 2.9-10.0 | 2.9-10.0 | 2.9-10.0 |
| Local resolution FSC threshold | 0.5 | 0.5 | 0.5 | 0.5 | 0.5 | 0.5 | 0.5 | 0.5 |
| **Refinement** |  |  |  |  |  |  |  |  |
| Initial model used (PDB code) | Alphafold3<br>prediction | Alphafold3<br>prediction | Alphafold3<br>prediction | Alphafold3<br>prediction | Alphafold3<br>prediction | Alphafold3<br>prediction |  |  |
| Model resolution (Å) | 3.6 | 3.5 | 3.7 | 3.3 | 3.2 | 3.2 |  |  |
| FSC threshold | 0.143 | 0.143 | 0.143 | 0.143 | 0.143 | 0.143 |  |  |
| Model composition |  |  |  |  |  |  |  |  |
| Non-hydrogen atoms | 80,980 | 70,541 | 44,040 | 33,096 | 52,070 | 32,688 |  |  |
| Protein residues | 10,543 | 9,617 | 5,730 | 4,125 | 6,545 | 4,170 |  |  |
| *B* factors (Å<sup>2</sup>) |  |  |  |  |  |  |  |  |
| Protein | 113.24 | 103.18 | 40.51 | 73.07 | 129.34 | 58.64 |  |  |
| R.m.s. deviations |  |  |  |  |  |  |  |  |
| Bond lengths (Å) | 0.003 | 0.002 | 0.003 | 0.005 | 0.004 | 0.004 |  |  |
| Bond angles (°) | 0.613 | 0.506 | 0.539 | 0.591 | 0.612 | 0.489 |  |  |
| Validation |  |  |  |  |  |  |  |  |
| MolProbity score | 2.11 | 1.92 | 1.47 | 1.63 | 1.71 | 1.53 |  |  |
| Clashscore | 10.96 | 7.94 | 6.85 | 5.30 | 7.12 | 4.25 |  |  |
| Poor rotamers (%) | 0.00 | 0.00 | 0.00 | 0.00 | 0.02 | 0.00 |  |  |
| Ramachandran plot |  |  |  |  |  |  |  |  |
| Outliers | 0.04 | 0.00 | 0.00 | 0.00 | 0.00 | 0.00 |  |  |
| Allowed | 10.02 | 7.88 | 2.47 | 4.95 | 4.58 | 4.67 |  |  |
| Favored | 89.84 | 92.12 | 97.53 | 95.05 | 95.42 | 95.33 |  |  |
| EMRinger score | 1.55 | 1.43 | 2.74 | 3.11 | 2.95 | 2.77 |  |  |
| Model vs. Data |  |  |  |  |  |  |  |  |
| CC(mask) | 0.79 | 0.79 | 0.83 | 0.86 | 0.80 | 0.87 |  |  |
| CC(volume) | 0.79 | 0.79 | 0.81 | 0.84 | 0.80 | 0.84 |  |  |

**Extended Data Table 2:** Modelled virion proteins of SWU1.

| Gene | Protein name | # Amino acids | Modelled | # Copies per phage | Symmetry |
|---|---|---|---|---|---|
| 67 | Tail tip | 548 | Partial | 3 | C3 |
| 70 | Tail spike | 604 | Yes | 3 | C3 |
| 71 | Tail hub | 336 | Yes | 6 | C6 |
| 72 | Tape measure | 837 | No | N/A | N/A |
| 75 | Tail tube | 198 | Yes | ~168 (6 per ring) | C6 |
| 76 | Tail terminator | 139 | Yes | 6 | C6 |
| 78 | Head-to-tail stopper | 122 | Yes | 6 | C6 |
| 79 | Head-to-tail adaptor | 125 | Yes | 12 | C12 |
| 81 | Capsid | 321 | Yes | 415 | I/C1 |
| 84 | Portal | 486 | Yes | 12 | C12/C1 |
| 89 | Tail fiber | 313 | Partial | 13 | C13/C1 |

